# Investigating Alzheimer’s Disease Biomarkers by Applying Machine Learning Models

**DOI:** 10.1101/2025.03.19.643368

**Authors:** Babak Khorsand, Shirin Salehi, Soroush Karimi, Sonia Karimipasand, Neda Fariborzi, Hamidreza Houri, Nastaran Asri

## Abstract

**Objective:** Alzheimer’s Disease (AD) is a debilitating neurodegenerative disorder characterized by memory loss, cognitive decline, and the accumulation of amyloid plaques and neurofibrillary tangles. This study investigates the interplay of various biomarkers and clinical features in diagnosing AD using machine learning (ML) techniques.

**Methods:** We analyzed data from 191 AD patients and 59 non-AD subjects, employing classifiers including Naive Bayes (NB), Random Forest (RF), Decision Tree (DT), Support Vector Machine (SVM), and K-Nearest Neighbors (KNN).

**Results:** Our findings indicate that KNN, SVM, RF, and DT achieved high sensitivity (94%) and accuracy (92%), demonstrating their potential as effective diagnostic tools. Notably, significant differences in feature values between AD patients and non-AD subjects suggest that biomarker-driven approaches can enhance diagnostic precision. Key biomarkers such as neprilysin, alpha-secretase, beta-secretase, amyloid plaques and urinary formic acid emerged as critical elements.

**Conclusion:** Our results underscore the importance of selecting a targeted subset of features to streamline the diagnostic process, allowing for more efficient and cost-effective screening. While our study reveals valuable insights into AD pathology and diagnosis, future research with larger, longitudinal cohorts is essential to further elucidate these relationships and enhance our understanding of Alzheimer’s mechanisms, ultimately aiming for innovative therapeutic strategies.

## Introduction

Alzheimer’s disease (AD) is a neurodegenerative disorder linked to memory loss and dementia. Its hallmark features include the presence of amyloid plaques, neurofibrillary tangles, inflammation, and a reduction in synapses. Age is a significant risk factor for developing Alzheimer’s, with women being more commonly affected than men. Research suggests that factors such as estrogen deficiency, hormonal changes, stress, and lifestyle may contribute to the higher prevalence in women [1]. The formation of extracellular amyloid-beta (Aβ) plaques and intracellular tau protein tangles is crucial to the disease’s development [2].

In a healthy brain, the Aβ peptide exists in a soluble form and is found at lower levels in the axons and synapses [3]. Normally, the clearance of Aβ from the brain significantly exceeds its production. Alzheimer’s disease may arise from an imbalance between the synthesis and degradation of amyloid, resulting in Aβ accumulation in the central nervous system. The hydrophobic properties of Aβ peptides, particularly the 42 variant, enable them to self-aggregate into various structures, ranging from dimers to low-molecular-weight oligomers, protofibrils, and fibrils. Ultimately, these accumulated filaments can lead to the formation of amyloid plaques [4].

Neprilysin (NEP) is a membrane-bound glycoprotein classified as part of the zinc metalloendopeptidase family. It plays a critical role in degrading Aβ, effectively breaking down both monomeric and oligomeric forms [5]. As individuals age, NEP levels decline, leading to an accumulation of Aβ. Another enzyme in this category is the insulin-degrading enzyme (IDE), which can break down multiple substrates that exhibit amyloidogenic characteristics. Reduced activity of IDE has been observed in patients with AD [6, 7].

Various gene mutations, including those in the amyloid precursor protein (APP), are linked to the development of AD [8, 9]. APP is a class I transmembrane protein, that undergoes processing by three key enzymes: α-secretase, β-secretase, and γ-secretase, which each play roles in non-amyloidogenic and amyloidogenic pathways. In the non-amyloidogenic pathway, APP is cleaved by α-secretase followed by γ-secretase, resulting in the production of 3-kDa peptide, termed p3, and an APP intracellular domain (AICD) fragments. Conversely, in the amyloidogenic pathway, β-secretase first cleaves APP, creating soluble amyloid precursor protein Beta (sAPPβ), which is then further processed by γ-secretase to produce AICD and amyloid-bAβ) peptide [10].

The timely and precise diagnosis of AD poses significant challenges in clinical practice. While PET scans and cerebrospinal fluid (CSF) biomarkers are commonly utilized in clinical research for diagnosis, they face limitations such as high costs and accessibility issues. In contrast, emerging blood-based markers offer promising potential for accurate, cost-effective, and widely available diagnostic tools, enabling earlier detection of the disease [11]. Recent studies have also explored the use of urine tests to assess formic acid levels as a novel diagnostic approach for AD. Formic acid, a byproduct of metabolizing formaldehyde, is typically low in healthy individuals, despite formaldehyde’s potential toxicity at elevated levels. Research suggests that while internal formaldehyde may contribute to cognitive functions like learning and memory, excessive concentrations in the brain can lead to cognitive impairments. Increased formic acid in urine may serve as an early warning indicator of AD [12].

Unraveling the critical elements that contribute to the progression of AD is essential in the quest for a cure. The advent of artificial intelligence (AI) and its various branches has significantly enhanced our ability to diagnose Alzheimer’s through innovative methods. In this article, we explore a range of characteristics to identify the most effective ones, employing machine learning (ML) techniques to classify AD. We applied input and output features across five classifiers: Naive Bayes (NB), Random Forest (RF), Decision Tree (DT), Support Vector Machine (SVM), and k-Nearest Neighbors (KNN). Ultimately, we assessed their performance using a confusion matrix, illuminating the path toward improved diagnostic accuracy.

## Material and Method

### Patients

This retrospective multicenter study included 191 AD patients (115 females and 76 males) from April 2022 to April 2023. AD diagnoses followed the National Institute on Aging— Alzheimer’s Association (NIA-AA) criteria [13]. Additionally, a control group of 59 patients (20 females and 39 males) with neurological conditions but without dementia was analyzed as non-AD subjects. The average age of the non-AD group was 76.17 ± 6.86 years, while AD patients had an average age of 77.14 ± 9.12 years. Relevant demographic data, laboratory results, and clinical characteristics for all participants were collected for evaluation.

### Features

In our study, we conducted a comprehensive analysis of a broad spectrum of demographic, clinical, and laboratory features, alongside relevant biomarkers. We assessed parameters including demographics (sex and age), cognitive function measured by the Mini-Mental State Examination (MMSE) to evaluate memory loss, and behavioral changes assessed through the Neuropsychiatric Inventory (NPI). Furthermore, we examined retinal changes via optical coherence tomography (OCT) to detect retinal thinning, and we analyzed movement disorders using electromyography (EMG). Additional evaluations included speech difficulties and dysphagia diagnosed through fiber-optic endoscopic evaluation of swallowing (FEES), as well as monitoring for weight loss. We documented neurological symptoms such as seizures, urinary incontinence (UI), and fecal incontinence (FI), and assessed metabolic markers including urinary formic acid levels and diabetes status through HbA1c and fasting blood sugar tests. Finally, we evaluated biomarkers such as amyloid plaques, alpha-secretase, beta-secretase, neprilysin, insulin-degrading enzyme (IDE), and dyshomeostasis of metal ions. This multifaceted approach enabled us to capture a detailed clinical profile and identify potential correlations among the various parameters studied.

### Classification Methods

We employed five ML classification methods including DT, SVM, NB, KNN, and RF [14]. DT recursively partitions data, providing interpretability but facing issues like overfitting. SVM establishes hyperplanes to separate classes, excelling with high-dimensional data yet struggling with large datasets and kernel selection. NB relies on conditional probability, offering simplicity and good performance with limited data, but often outperformed by more complex models. KNN uses distance-based decision-making, proving effective for classification and anomaly detection. RF builds multiple decision trees and aggregates results, mitigating overfitting and handling large datasets effectively despite computational costs [15-20].

### Evaluation measures

The following measures are used for evaluating our models:

Sensitivity is a widely used metric derived from the matrix. It represents the proportion of patients with the disease who receive a positive test result, along with data regarding individuals without the disease.

Specificity refers to the percentage of patients who do not have the disease and receive a negative test result, providing no information about those who do have the disease.

Accuracy is the most straightforward and widely used metric derived from the confusion matrix. It reflects the classifier’s ability to identify both positive and negative classes correctly.

Another important metric called the F-Measure is commonly used to evaluate classification performance. It represents the harmonic mean of precision (positive predictive value) and recall (sensitivity).

### Important Features

To extract important features, we eliminate each feature one at a time and reconstruct the models without that feature. We then measure the decrease in accuracy of the new models compared to the original models. The amount of accuracy lost is considered the score of each feature. Variables with the highest decrease in accuracy are deemed the most important. Afterward, we use only the most important features to reconstruct the models and evaluate their effectiveness.

## Result

In the methodology section, we outlined our analysis approach, which involved a variety of features. Figure 1 illustrates these features on the horizontal axis, with their values represented as a heatmap on the vertical axis. Our classification framework is binary, and feature values have been normalized to a scale of 0 to 1. A bold purple (value: 1) indicates the presence of a feature, while a pale purple (value: 0) signifies its absence. The class column on the far right reveals a distribution of 76.4% bold purple (patients diagnosed with AD) and 23.6% pale purple (patients without AD).

**Figure 1.**
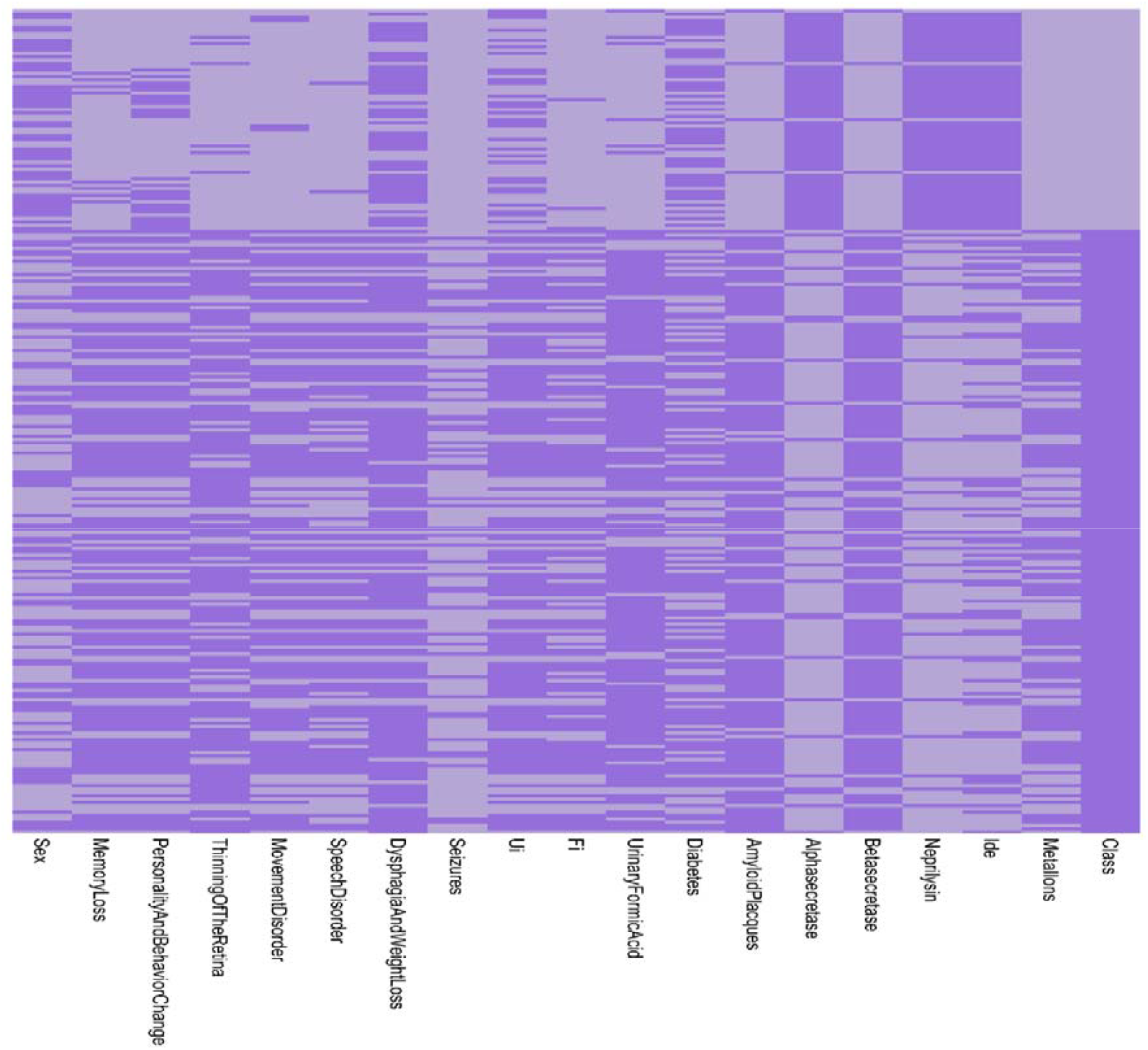
Heatmap illustrating all features and their corresponding values. In each column, bold purple indicates the presence of the related feature, while pale purple indicates its absence. The rightmost column represents the class distribution, with bold purple denoting patients with AD and pale purple denoting patients without AD.

**Figure 2:**
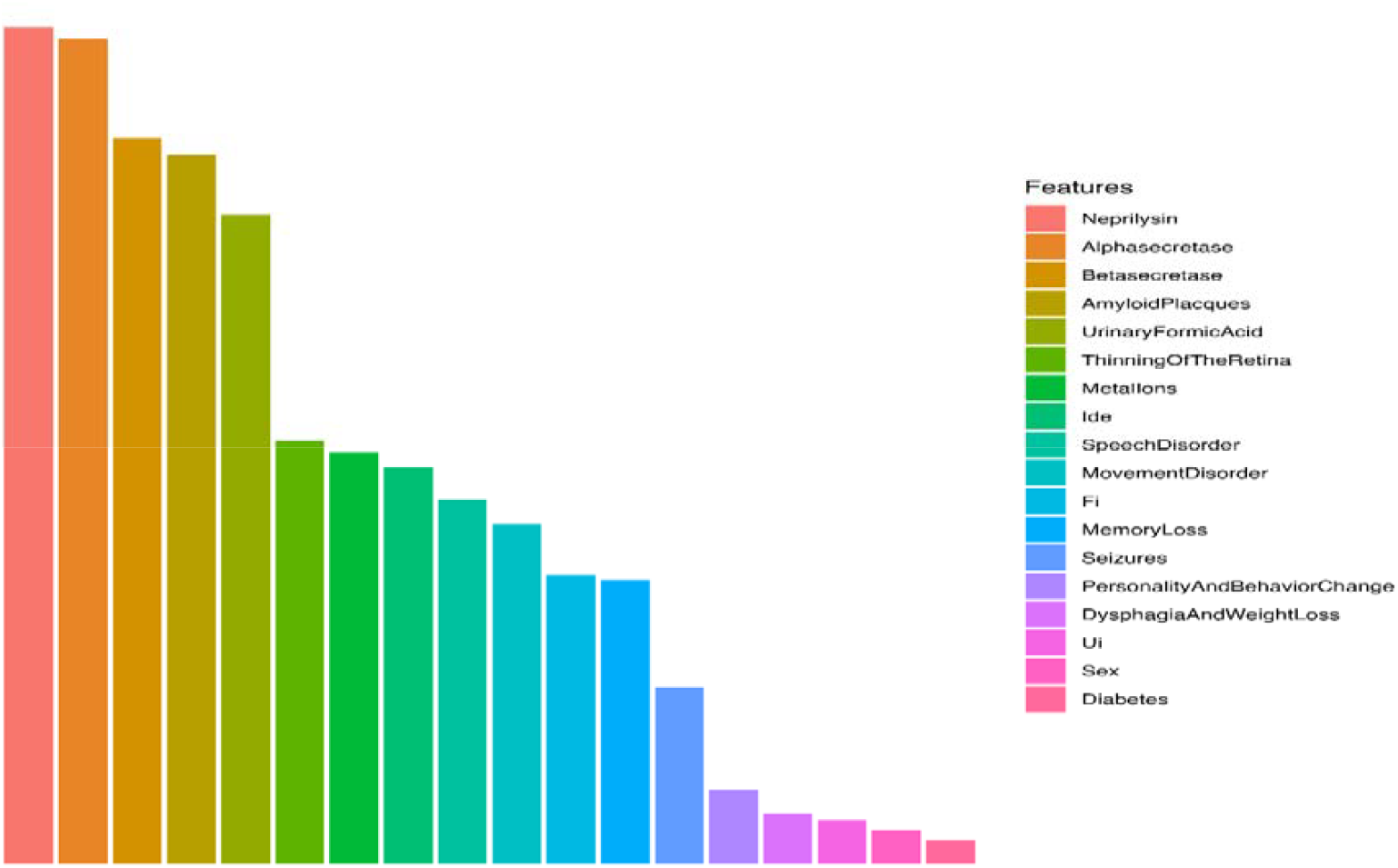
The outcomes of diverse models (DT, KNN, NB, RF, and SVM) across multiple measures including sensitivity, specificity, F-measure, and accuracy.

Our model was developed using five advanced algorithms including DT, KNN, NB, RF, and SVM. We assessed their performance through several metrics, including specificity, accuracy, F-measure, and sensitivity.

In terms of specificity, the models achieved the following scores: KNN (84%), NB (92%), SVM (84%), DT (84%), and RF (84%). For accuracy, the results were impressive: KNN (92%), NB (90%), SVM (92%), DT (92%), and RF (92%). When evaluating sensitivity, we found the following: KNN (94%), NB (89%), SVM (94%), DT (94%), and RF (94%). Finally, regarding the F-measure, the models demonstrated: KNN (94%), NB (89%), SVM (94%), DT (94%), and RF (94%).

In our variable analysis, we identified several important features that significantly distinguished AD patients from non-AD individuals (Figure 3). Our results highlight notable differences in various features, including neprilysin (score= 2.89), alphasecretase (score= 2.86), betasecretase (score= 2.51), amyloid plaques (score= 2.45), and urine formic acid (score= 2.25). These findings emphasize the relevance of these parameters in the clinical evaluation and diagnosis of AD.

**Figure 3:**
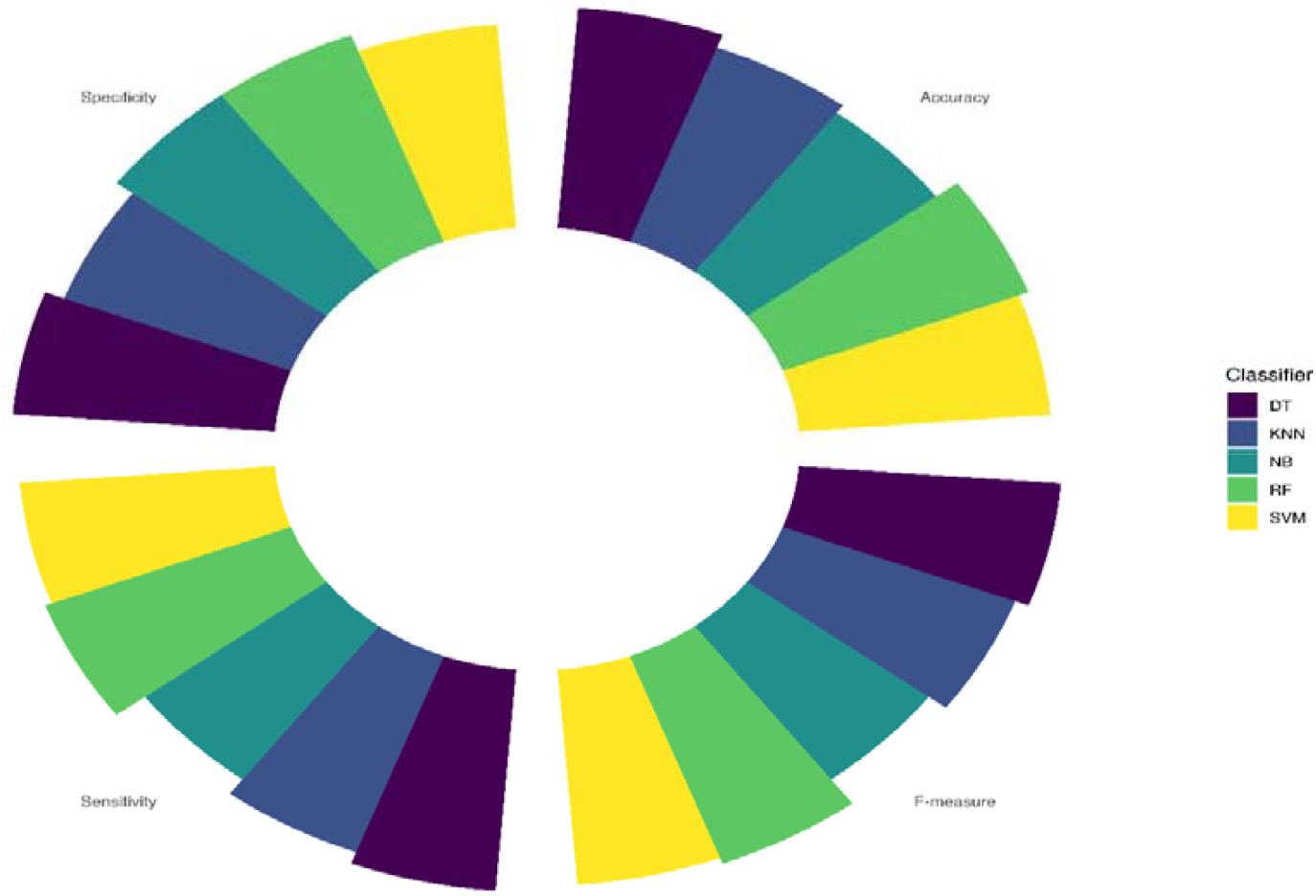
All features were sorted from most main feature to lowest main feature and the five tops were selected for the next studies to unravel their importance.

To evaluate the effectiveness of these key features, we employed a visually engaging Radar chart. This chart showcased the performance of various models, including DT, RF, SVM, KNN, and NB, while highlighting crucial evaluation metrics such as specificity, sensitivity, and accuracy. Our findings demonstrated remarkably consistent results, with no noticeable differences between the approaches. This indicates that leveraging these essential features proved to be both time-efficient and cost-effective for AD diagnosis (Figure 4)

**Figure 4.**
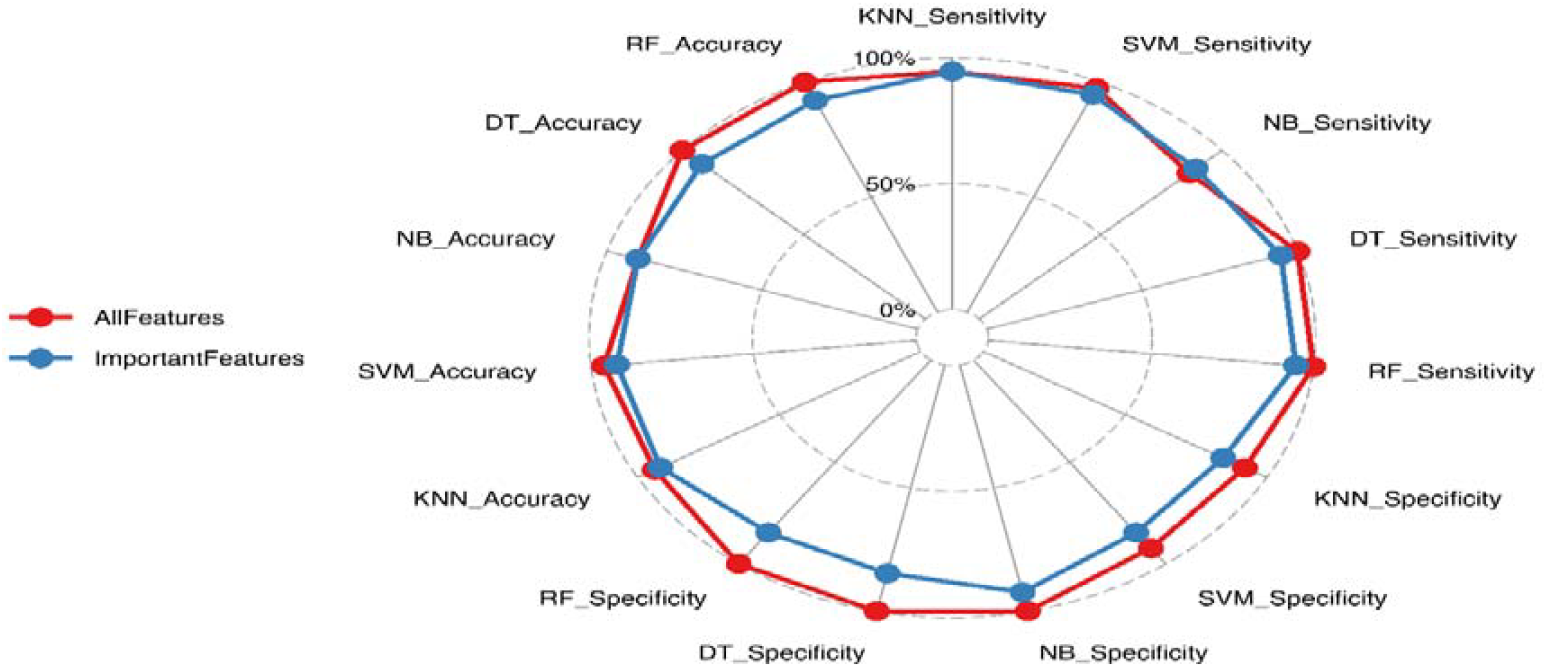
The elucidation of both essential features and the entire feature set significance through a Radar chart, employing DT, RF, SVM, KNN and NB models focusing on sensitivity, specificity, and accuracy. The data depict not any noticeable changes in models’ functions.

## Discussion

AD is a progressive brain disorder that leads to irreversible damage, characterized by key pathological features such as amyloid plaques, which are vital for its diagnosis and understanding [21]. This condition causes permanent damage to memory and cognitive function, ultimately resulting in death. Therefore, early detection is crucial for enhancing patient care and advancing clinical research [21]. As one of the most costly diseases to treat, Alzheimer’s has garnered significant interest from researchers aiming to develop highly accurate automated algorithms for early diagnosis. However, identifying and predicting Alzheimer’s in its initial stages can be quite challenging [21].

In recent years, the use of computer-aided analysis models powered by ML and AI has significantly improved the examination of various neuroimaging techniques and non-imaging biomarkers (25). Research has increasingly applied ML to forecast the progression of AD. For instance, a study developed a hybrid ML framework to analyze longitudinal data for predicting dementia outcomes in patients with Mild Cognitive Impairment (MCI). Although this model achieved an impressive accuracy of 87.5% using Random Forest, it showed variability across different performance metrics and a notable sensitivity of 92.9%, which affected its specificity, recorded at only 58.3% (26). Another study identified 15 clinical variables associated with MCI progression, achieving accuracy, sensitivity, and specificity rates of 71%, 67.7%, and 71.7%, respectively (26).

In this article, we explored a range of AD related clinical features and biomarkers using five different classifiers. Our findings highlight that KNN, SVM, RF, and DT performed similarly well, affirming their potential as effective diagnostic tools. In accordance with our finding, Alalayah et al. [22] demonstrated that RF, MLP, DT SVM and KNN achieved accurate and effective diagnostic results for diagnosing Parkinson’s disease (PD) as a neurodegenerative disorder. Dara et al. [23] study reviews over 80 publications since 2017 on AD detection using ML techniques like SVM, DT, and ensemble models. It examines 50 papers with specific architectural approaches. The literature is categorized into data-related, methodology-related, and medical-related components to highlight challenges in the field. The conclusion discusses future research opportunities, recommending a focus on feature selection and optimization techniques (e.g., whale and gray wolf optimization) to enhance diagnostic accuracy, particularly using MRI images.

Research indicates that analyzing specific biomarkers helps identify preclinical stages of diseases and supports the development of new targets for disease-modifying therapies [24]. The significant variations in feature values between AD patients and non-AD subjects in the current study also suggest that biomarker-driven methods could further refine diagnostic precision. Importantly, NEP and secretase activities emerged as pivotal factors, underscoring the essential role of Aβ metabolism and clearance in the pathology of AD. These findings align with previous studies showing that reduced NEP activity is linked to increased Aβ accumulation, reaffirming its potential as a therapeutic target [6, 7]. Furthermore, our analysis reveals that other parameters, such as urinary formic acid could offer additional, clinically relevant insights into AD progression, warranting further exploration.

In a study conducted by Saad Alatrany et al. [25] Using a dataset of 169,408 records, SVM models demonstrated high performance, achieving an F1 score of 98.9% in binary classification and 88% in predicting AD progression. To enhance understanding, rule-extraction methods highlighted key factors like MEMORY and JUDGMENT, validated by SHAP and LIME models. Chang et al. [26] in their study highlighted the potential of ML combined with novel biomarkers for improving AD diagnosis. They introduced key biomarkers including neurofilament light (NFL) for neuronal injury, neurogranin, beta-site amyloid precursor protein cleaving enzyme (BACE)1, synaptotagmin, synaptosome associated protein 25 (SNAP-25), Growth-associated protein 43 (GAP-43), and synaptophysin for synaptic dysfunction, and soluble triggering receptor expressed on myeloid cells 2 (sTREM2) and chitinase-like protein YKL-40 for neuroinflammation. They stated that traditional methods are costly and invasive, but using ML algorithms can enhance diagnostic accuracy.

Interestingly, our study revealed that focusing on a subset of essential features yielded nearly identical performance metrics compared to using the full feature set. This insight suggests that targeted biomarker evaluations could streamline the diagnostic process, reducing costs and time while maintaining accuracy. Prioritizing effective biomarkers such as NEP, alphasecretase, amyloid plaques and urinary formic acid may thus facilitate more efficient screening and monitoring strategies in clinical practice.

While our study provides valuable insights, it is important to acknowledge its limitations, including the retrospective nature and the relatively small sample size. Future research should aim to replicate these findings in larger cohorts and explore longitudinal data to assess how features evolve over time in AD progression. Additionally, integrating advanced AI algorithms may yield enhanced predictive capabilities and facilitate the discovery of new biomarkers, further advancing our understanding of this intricate disease.

In conclusion, our study highlights the critical interplay between key biomarkers and the pathogenesis of AD, reinforcing the significance of neprilysin, amyloid plaques, alphasecretase, betasecretase, and urinary formic acid in understanding and diagnosing this complex neurodegenerative disorder. The strong performance of machine learning classifiers—particularly KNN, SVM, Random Forest, and Decision Tree—demonstrates that integrating these biomarkers can lead to improved diagnostic accuracy and sensitivity.

By elucidating the relationships between these biomarkers, we enhance our understanding of AD mechanisms, offering valuable insights for possible therapeutic targets. Additionally, the findings advocate for streamlined screening practices that prioritize these essential features, ensuring efficient and cost-effective diagnostic protocols. Future research should focus on expanding the cohort size and utilizing longitudinal studies to track biomarker evolution over time. As we move forward, combining biological insights with advanced computational methods will be crucial in our pursuit of effective treatments and ultimately, a cure for Alzheimer’s Disease. This multidisciplinary approach has the potential to transform how we diagnose and manage AD, enriching the lives of those affected by this debilitating condition.

## Disclosure of interest

The authors report there are no competing interests to declare

## Ethics approval

The study was approved by the ethical committee of the Shahid Beheshti University of Medical Sciences in Tehran, Iran (IR.SBMU.RETECH.REC.1403.684).

## Notes

### Competing Interest Statement

The authors have declared no competing interest.

